# Unconventional Activation of IRE1 Enhances TH17 Responses and Promotes Neutrophilic Airway Inflammation

**DOI:** 10.1101/2023.06.30.547286

**Authors:** Dandan Wu, Xing Zhang, Kourtney M. Zimmerly, Ruoning Wang, Amanda Livingston, Takao Iwawaki, Ashok Kumar, Xiang Wu, Michael A. Mandell, Meilian Liu, Xuexian O. Yang

## Abstract

Treatment-refractory severe asthma manifests a neutrophilic phenotype associated with TH17 responses. Heightened unfolded protein responses (UPRs) are associated with the risk of asthma, including severe asthma. However, how UPRs participate in the deregulation of TH17 cells leading to this type of asthma remains elusive. In this study, we investigated the role of the UPR sensor IRE1 in TH17 cell function and neutrophilic airway inflammation. We found that IRE1 is induced in fungal asthma and is highly expressed in TH17 cells relative to naïve CD4^+^ T cells. Cytokine (e.g. IL-23) signals induce the IRE1-XBP1s axis in a JAK2-dependent manner. This noncanonical activation of the IRE1-XBP1s pathway promotes UPRs and cytokine secretion by TH17 cells. *Ern1* (encoding IRE1)-deficiency decreases the expression of ER stress factors and impairs the differentiation and cytokine secretion of TH17 cells. Genetic ablation of *Ern1* leads to alleviated TH17 responses and airway neutrophilia in a *Candida albicans* asthma model. Consistently, IL-23 activates the JAK2-IRE1-XBP1s pathway *in vivo* and enhances TH17 responses and neutrophilic infiltration into the airway. Taken together, our data indicate that IRE1, noncanonically activated by cytokine signals, promotes neutrophilic airway inflammation through the UPR-mediated secretory function of TH17 cells. The findings provide a novel insight into the fundamental understanding of IRE1 in TH17-biased TH2-low asthma.

## Introduction

Asthma, one of the most common chronic diseases, is characterized by uncontrolled lung inflammation that constricts the airways, leading to breathing difficulty. Asthma affects about 262 million people and caused 461,000 deaths in 2019 (World Health Organization data). Among those, about half of the cases are T helper 2 (TH2 or T2)-biased ^1^, whereas non-eosinophilic T2-low endotype is more common in patients with severe asthma ^2, 3^, and is associated with a high risk of mortality and disability, causing a huge socio-economic burden. T2-low asthma, generally characterized by neutrophilic or paucigranulocytic inflammation, is often refractory to corticosteroid treatment even combined with bronchodilators ^4–6^.

Neutrophilic severe asthma is closely associated with TH17 responses ^4, 5, 7–9^. TH17 cells play an important role in immunity against extracellular pathogens and regulate tissue inflammation by expressing pro-inflammatory cytokines IL-17 (or IL-17A), IL-17F, and IL-22 ^10, 11^. Airway colonization of bacteria and fungi induces TH17 responses ^12–14^, and leads to airway neutrophilia ^13, 15^. Compared with healthy controls and those with mild to moderate asthma, patients with severe asthma have increased numbers of IL-17^+^ and IL-17F^+^ cells in their bronchial biopsies ^16^, although there is no significant increase in the levels of IL-17 in the sera or sputum ^17, 18^, suggesting an airway localized TH17 response. Transcriptomic analysis demonstrated a strong upregulation of IL-17-inducible chemokines CXCL1, 2, 3, and 8 (IL-8), and cytokine CSF3 (G-CSF), IL-22-inducible chemokine CCL3, and TH17 cell-expressed chemokines CCL3 and Galectin-3 in airway brushings from patients with severe neutrophilic asthma relative to those of control subjects ^19–21^. Among these mediators, CSF3 promotes granulopoiesis and regulates neutrophil function ^22^, and CXCL8, CCL3, and Galectin-3 are neutrophil chemoattractants, which together govern neutrophilic airway influx. In addition, we and others have shown that IL-17 induces matrix metalloproteinases (MMPs) 3, 9, 12, and 13 ^23–25^, mediating tissue remodelling. In this context, the MMP12 serine 357 variant is linked with clinically worse asthma with an increased risk of exacerbations ^26^. Besides TH17 cells, group 3 innate lymphocytes (ILC3s), a subset of TH17-type lymphocytes, are implicated in obesity-associated airway hyperresponsiveness (AHR) in mice ^27, 28^. Therefore, deregulation of TH17-type responses is essential in neutrophilic airway inflammation.

The unfolded protein response (UPR) is a cellular homeostatic adaptation to endoplasmic reticulum (ER) stress caused by massive protein synthesis. The UPR is initiated by three major ER transmembrane-associated sensors: inositol-requiring enzyme 1 (IRE1), protein kinase RNA-like ER kinase (PERK), and activating transcription factor 6 (ATF6). Among these, ATF6 and IRE1, are the effector pathways promoting cell secretory function. After activation, ATF6 and IRE1 regulate *Xbp1* gene (encoding unspliced *Xbp1u* mRNA) transcription and splicing (to functional *Xbp1s* mRNA), respectively ^29–31^. XBP1s is essential in the secretory function of cells, including immune cells ^29, 30^. GWAS studies have associated the ER stress factor Orosomucoid-like (ORMDL) 3 with the risk of asthma ^32–34^, including severe asthma ^35, 36^. ORMDL3 promotes airway smooth muscle hyperplasia *in vitro* ^37^. In an *Alternaria* asthma model, ORMDL3 promotes T2-high airway disease through activation of the ATF6 arm of UPR in airway epithelial cells ^38^. Human airway epithelial brushings from asthmatic subjects also demonstrate an upregulation of several ATF6-related transcripts compared with those from healthy controls ^39^. Despite that in airway structural cells, little is known about the role of UPRs in T helper (TH) cells.

TH cells (and also immune infiltrates) belong to secretory cells. When receiving stimuli from T cell receptor (TCR) engagement, costimulation, and cytokine signals, TH cells produce copious amounts of effector molecules, including their signature cytokines, such as TH17 cytokines (IL-17, IL-17F, IL-22, and GM-CSF). During this process, the accumulation of misfolded/unfolded proteins in the ER lumen leads to ER stress and subsequent UPRs. The IRE1a-XBP1 pathway has been shown to regulate TH2 responses in airway allergy ^40, 41^. Recently, we have demonstrated that the UPR factor ATF6 is selectively expressed by TH2 and TH17 cells, in which pSTAT6 and pSTAT3, respectively drive ATF6 expression ^42^. ATF6 in turn promotes UPRs and boosts the expression of TH2 and TH17 cytokines that activate STAT6 and STAT3. This feedforward loop augments TH2 and TH17 responses, leading to enhanced mixed granulocytic airway inflammation in an experimental asthma model, in which T cell-specific *Atf6* deficiency or inhibition of ATF6 moderately lessens airway neutrophilia ^42^. It is unclear whether IRE1, another major effector UPR factor, also participates in the pathology of neutrophilic asthma through regulation of TH17 response. IRE1 has two isoforms, IRE1α (encoded by *Ern1*) and IRE1β (encoded by *Ern2*). The former is ubiquitously expressed by all types of mammalian cells, while the latter is restrictedly expressed by mucosal epithelial cells ^43^. In this study, we investigated the role of IRE1α (referred to IRE1, hereafter) in TH17 cells that drive neutrophilic fungal asthma. We found that IRE1 was induced by TH17 skewing cytokines, IL-23 and IL-6, and TCR-costimulatory signals. Previously, we and others demonstrated that cytokines leptin and IL-3 induce IRE1 activation in TH2 and pro-B cells, respectively ^41, 44^. How such extracellular signals activate IRE1 is entirely unknown. Because IL-23, IL-6, and leptin activate JAK2 ^45^ (leptin also activates MEK and mTOR), we asked whether JAK2 could activate IRE1 and found that JAK2, activated by IL-23, interacted with and directly phosphorylated IRE1. This activation of IRE1 by cytokine signals is distinct from the traditional view that unfolded or misfolded proteins-caused dissociation of the ER chaperon BiP from IRE1 leads to IRE1 activation ^46, 47^. The noncanonical activation of IRE1 promoted cytokine secretion by TH17 cells in an XBP1s-dependend manner. Genetic ablation of IRE1 in T cells resulted in alleviated TH17 development and airway neutrophilic infiltration in a *Candida albicans* asthma model. In this context, IL-23 treatment enhanced the IRE1-XPB1s axis and intensified TH17 response-associated airway neutrophilia. Taken together, our data suggest that the cytokine**−**IRE1**−**XBP1s axis upregulates the responsiveness and secretory function of pro-inflammatory TH17 cells, leading to airway neutrophilia. These findings provide a novel insight into the fundamental understanding of the role of UPRs in TH17 cell-mediated T2-low asthma.

## Results

### IRE1 is induced during neutrophilic asthma and promotes TH17 differentiation

Fungal airway infection elicits protective TH responses. For instance, *Candida albicans* elicits both TH17 and TH2 responses during airway infection in murine ^48^. However, uncontrolled fungal colonization in the airway has been associated with the development and exacerbation of human neutrophilic asthma through induction of TH17 responses ^13^, in which fungi (e.g., *C. albicans*)-induced IL-23 drives TH17 (and Vδ1 T) Cell responses ^49, 50^. *C. albicans* is a major fungal inducer of human TH17 responses ^51^. In a neutrophilic asthma model, we found that *Ern1* mRNA was induced by intranasal administration of heat-inactivated *C. albicans* (**Fig. 1A**). *Ern1* mRNA was also up-regulated in TH17 cells compared with naïve CD4^+^ T cells (**Fig. 1B**), suggesting that IRE1 may promote asthmatic reaction through control of TH17 responses. We then generated T cell-specific *Ern1*-deficient (*Ern1*^fl/fl;Cd4Cre^) mice by crossing *Ern1*^fl/fl^ with Cd4Cre mice ^52, 53^, and examined the role of IRE1 in TH17 differentiation. *Ern1*-deficiency led to decreased IL-17^+^ cell frequencies and IL-17 cytokine production (**Fig. 1C-D**). Consistently, *Ern1*-deficient TH17 cells expressed decreased levels of mRNAs of TH17 signature genes, *Il17a* (encoding IL-17), *Il17f*, *Rorc(gt)*, *Il6*, and *Il21*, which may be explained by the downregulation of the master transcription factor RORγt of TH17 cells, and IRE1 and XBP1s target genes, *Xbp1u*, *Xbp1*s, *Hspa5* (encoding BiP), and *Ddit3* (encoding CHOP) ^54–56^ (**Fig. 1E-F**). Therefore, IRE1 plays a vital role in TH17 differentiation and function and may promote neutrophilic inflammation in asthma.

**Figure 1.**
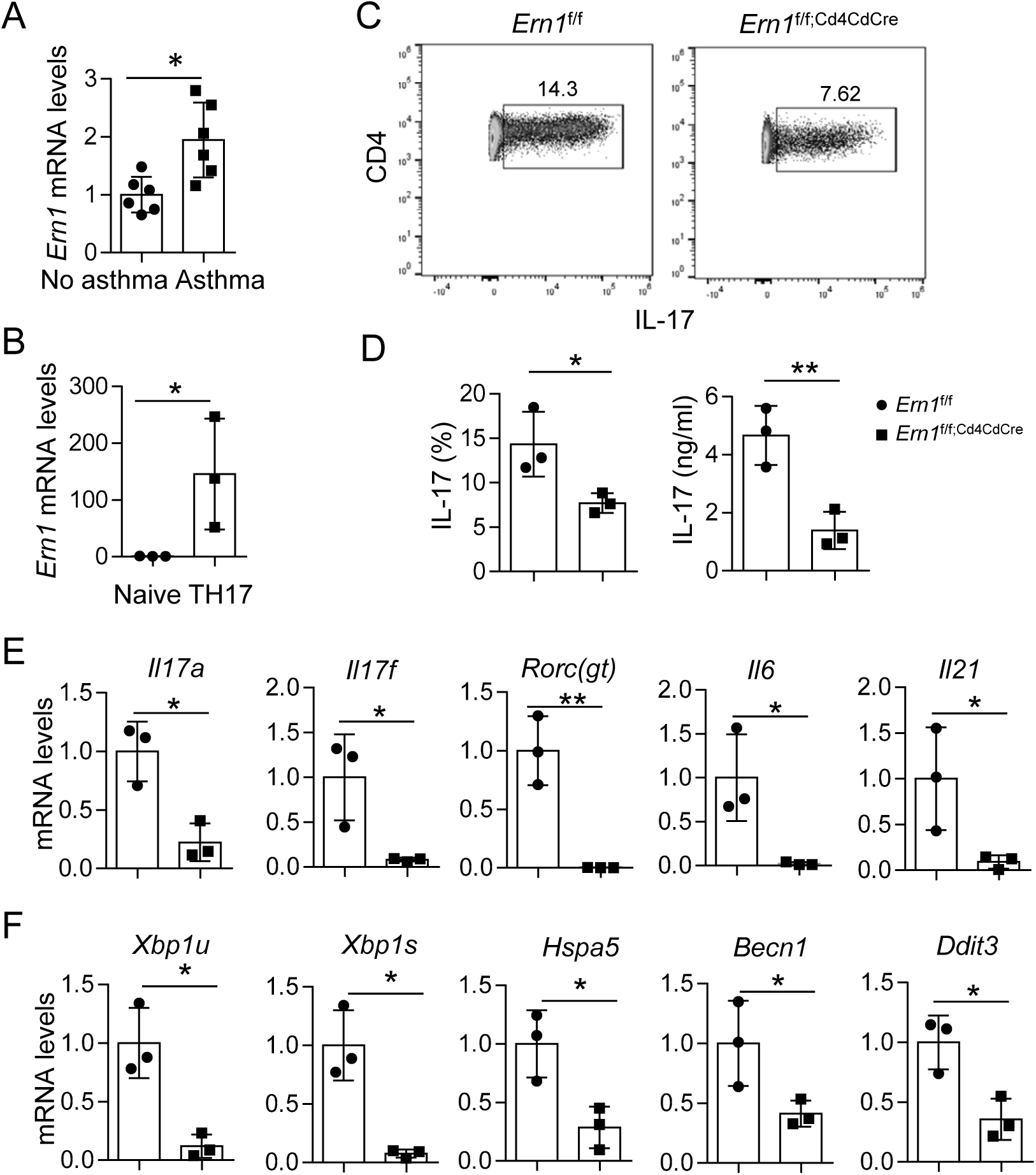
IRE1 is induced in neutrophilic asthma and intensifies TH17 differentiation and function. (A) RT-qPCR of *Ern1* mRNA in lungs from asthmatic and healthy mice. Asthma was elicited by using *Candida albican* extract. n = 6 per group. (B) RT-qPCR of *Ern1* mRNA in TH17 cells and naive CD4^+^ T cells. (C-F) Characterization of TH17 cells polarized from *Ern1*^fl/fl^ and *Ern1*^fl/fl;Cd4Cre^ naïve CD4^+^ T cells. (C) Flow cytometry of IL-17^+^ CD4^+^ T cells. (D) Statistical analysis of IL-17^+^ CD4^+^ T cell frequencies (left) and ELISA of IL-17 expression (rigt) in (C). (E) RT-qPCR of mRNA expression of TH17 signature genes, *Il17*, *Il17f*, *Rorc(gt)*, *Il6*, and *Il21*. (F) RT-qPCR of mRNA expression of IRE1 downstream genes, *Xbp1u*, *Xbp1s*, *Hspα5*, *Becn1*, and *Ddit3*. mRNA values were normalized to the internal reference gene *Actb* (A-B, E-F). Data (mean ± SD) shown are a representative (A, C) or a combination (B, D-F) of 3 experiments. Student’s *t*-test, \**p* < 0.05; ** *p* < 0.005.

### T cell-specific Ern1-deficiency alleviates neutrophilic airway inflammation

Since IRE1 is induced in asthmatic lungs and regulated UPRs and functions of TH17 cells (**Fig. 1**), we then tested the role of IRE1 in Candida-caused airway inflammation, driven by TH17 responses. *Ern1*^fl/fl^ and *Ern1*^fl/fl;Cd4Cre^ mice were intranasally (i.n.) immunized with *C. albicans* extract plus a model antigen chicken ovalbumin (OVA). We found that levels of XBP1s in lung infiltrating TH17 cells from *Ern1*^fl/fl;Cd4Cre^ mice were significantly lower than those of control mice (**Fig. 2A**), suggesting that *Ern1*-deficiency impairs XBP1s expression. *Ern1*-deficiency in T cells resulted in decreased frequencies of TH17 (IL-17^+^) cells in the CD4^+^ fraction of bronchoalveolar lavage fluid (BALF) (**Fig. 2B**) and decreased levels of TH17 cytokines, IL-17 and IL-22, in BALF (**Fig. 2C**). Upon *ex vivo* recall with various concentrations of OVA, lung-draining mediastinal lymph node (LLN) cells from *Ern1*^fl/fl;Cd4Cre^ mice expressed lower amounts of TH17 cytokines than those from control mice (**Fig. 2D**). Consistently, *Ern1*^fl/fl;Cd4Cre^ mice had fewer airway influxes of macrophages and neutrophils compared with control mice (**Fig. 2E**). Hematoxylin and eosin (H&E) staining further revealed that lungs of *Ern1*^fl/fl;Cd4Cre^ mice displayed profound decreases immune infiltration including neutrophils at the peribronchial and perivascular spaces than those of *Ern1*^fl/fl^ mice (**Fig. 2F**). Collectively, the IRE1−XPB1s axis governs TH17 responses and neutrophilia in Candida-induced airway disease.

**Figure 2.**
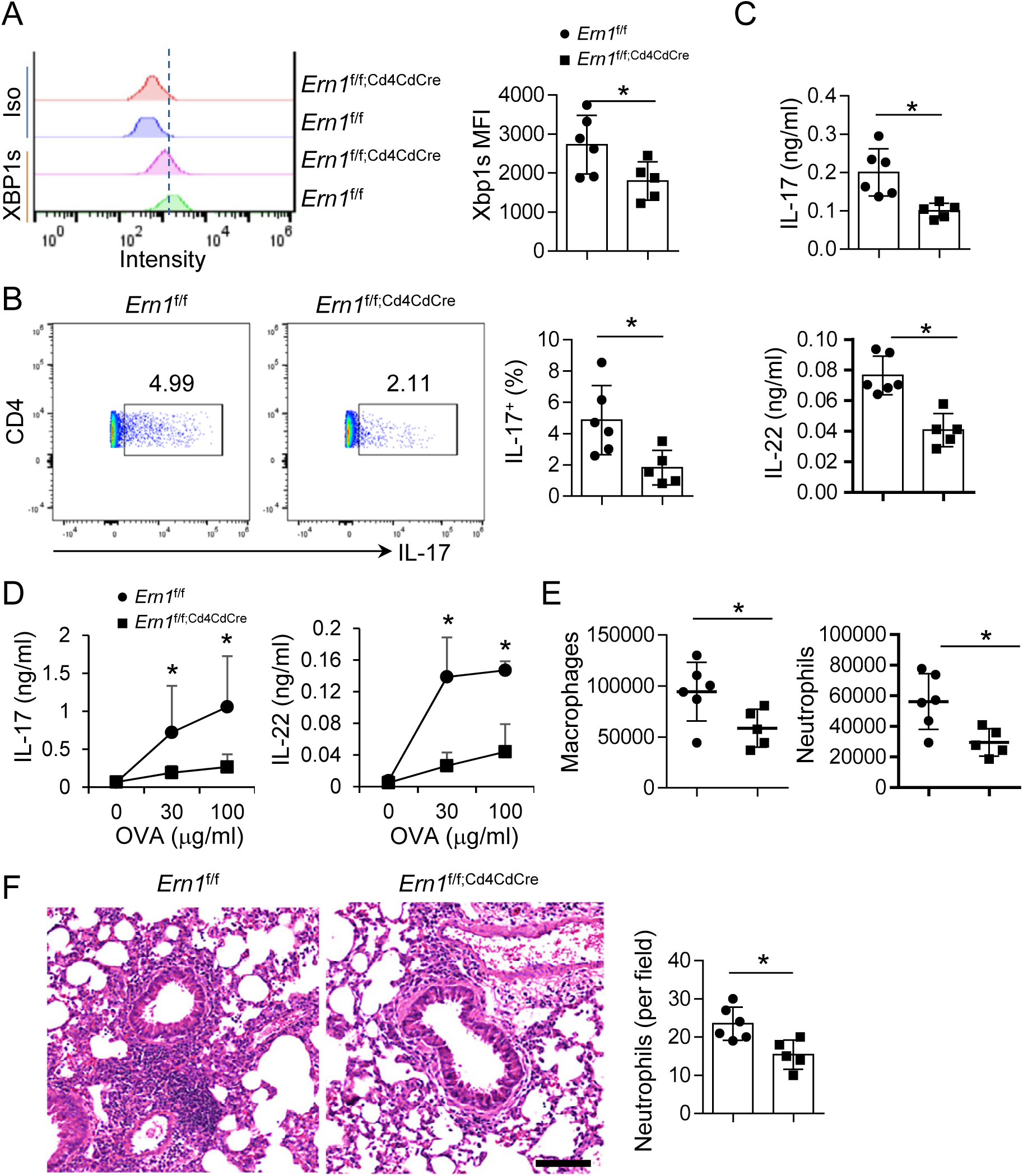
T cell-specific *Ern1*-deficiency alleviates neutrophilic airway inflammation. Neutrophilic asthma was induced in *Ern1*^fl/fl^ and *Ern1*^fl/fl;Cd4Cre^ mice by i.n. challenges with *C. albicans* extract and OVA. (A) Intracellular stain of XBP1s in asthmatic lung infiltrating TH17 cells on a CD3^+^ CD4^+^ IL-23R^+^ gate. Iso, isotype control antibody. (B) Intracellular stain of CD4^+^ IL-17^+^ cells in LLNs on a CD3^+^ gate. (C-D) ELISA of cytokines in BALF (C) and culture supernatant of LLNs after *ex vivo* recall with OVA (D). (E) Profiles of macrophages and neutrophils in BALF. (F) H&E stain of lung sections. Scale bar, 50µm. Statistical analysis of neutrophils (average counts per 20x field per sample). Data (mean ± SD) are a representative of two experiments. n = 5-6 per group. Student’s *t* test, \**p* < 0.05.

### Extracellular signals activate the IRE1-XBP1s axis in TH17 cells

During ER stress, activated ATF6 and IRE1 regulate XBP1s expression through transactivation of the Xbp1 gene and splicing Xbp1u mRNA to Xbp1s, respectively ^29–31^. We previously showed that leptin induces XBP1s expression through activation of IRE1 in TH2 cells ^41^. TH17 cells respond to many micro-environmental stimuli. To better understand whether and how these stimuli activate the IRE1 pathway in TH17 cells, we tested extracellular stimuli essential for TH17 differentiation and/or function. We found that both TCR-costimulation and cytokine signals from IL-23 and IL-6 activated the IRE1−XBP1s axis in TH17 cells (**Fig. 3A**). However, IL-23 and leptin did not alter ATF6 activation in TH17 and TH2 cells, respectively (ref. ^41^ and Supplemental **Fig. S1A**). These data indicate that cytokine signals activate IRE1 but not ATF6, leading to the induction of XBP1s.

**Figure 3.**
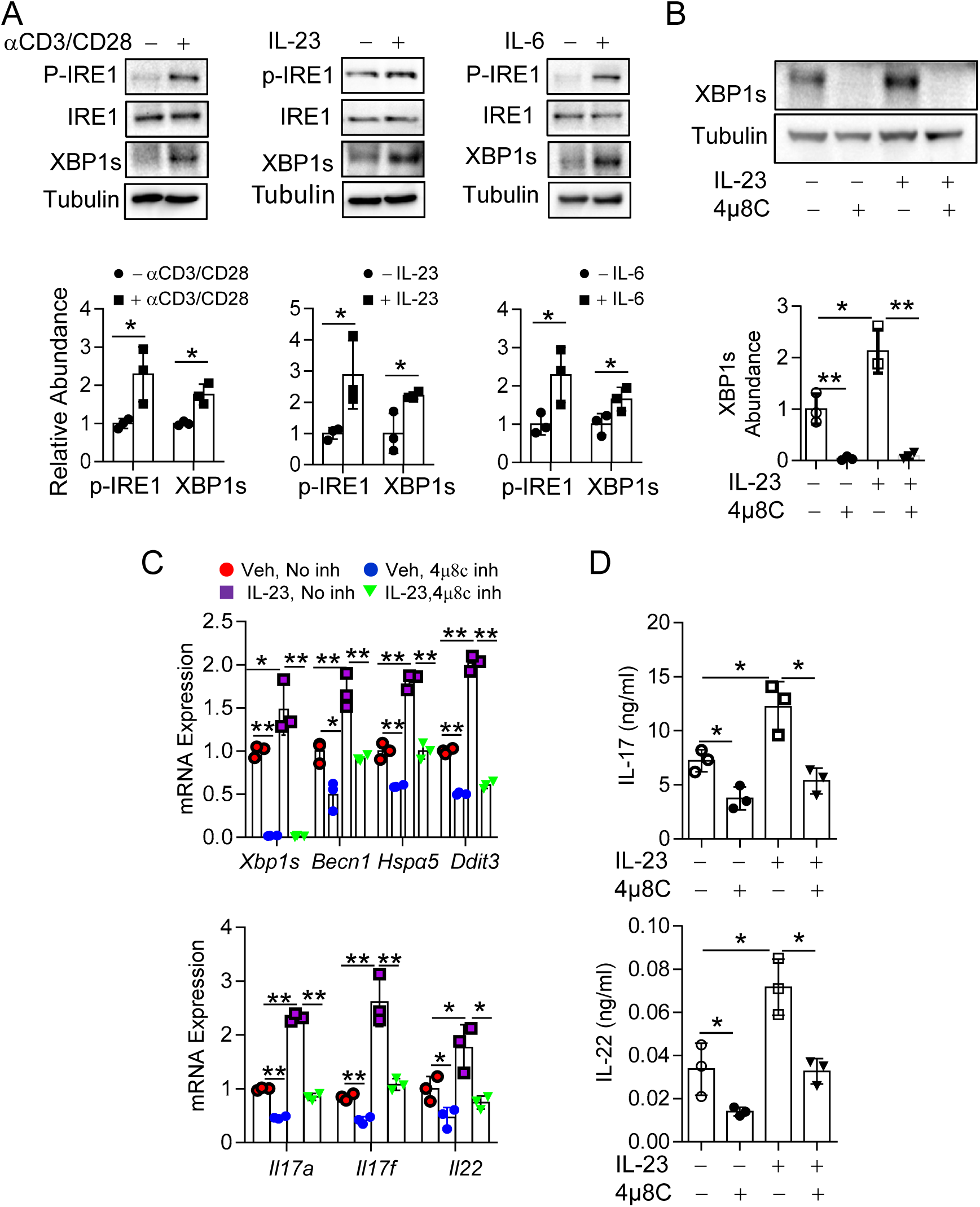
Extracellular signals activate the IRE1-XBP1s cascade and control TH17 function. (A) Western blot of p-IRE1 and XBP1s expression in TH17 cells treated with or without anti- CD3/CD28, IL-23, or IL-6. p-IRE1 and XBP1s abundances were relative to IRE1 and Tubulin, respectively. (B) Western blot XBP1s in TH17 cells treated with 10 µM IRE1 inhibitor 4μ8C or a vehicle in the presence or absence of IL-23. Abundances of XBP1s was normalized to Tubulin. (C) RT-qPCR of mRNA expression of indicated genes in TH17 cells treated as (B). *Actb* was used as a loading control. (D) ELISA of cytokines, IL-17 and IL-22. Data (means ± SD) were pooled from 3 independent experiments. Student’s *t* test, \**p* < 0.05, \*\**p* < 0.005.

### IL-23 induces XBP1s expression through activation of IRE1

Among the extracellular stimuli, we are interested in the cytokines that are important for TH17 cell function and maintenance, since manipulation of their signalling may allow us to impede the pathogenic function of these cells. IL-23 stimulates the development of pathogenic TH17 cells ^25, 57–59^. In the absence of IL-23-IL-23R signals, TH17 cells cannot fully undergo clonal expansion and egress from the draining lymph node to the bloodstream and tissues, and therefore, fail to mediate encephalitogenicity in experimental autoimmune encephalomyelitis (EAE, a mouse model of human MS) ^60^. In asthma, fungus-induced IL-23 contributes to the relapse and exacerbation of the disease through enhancing TH17 responses ^49, 50^. Since IL-23 does not activate ATF6 (Supplemental **Fig. S1A**), we asked whether IL-23 induces XBP1s expression through activation of IRE1 in TH17 cells. To address this, we used an IRE1 selective inhibitor 4μ8C ^61^. Inhibition of IRE1 resulted in decreased expression of XBP1s, and meanwhile, diminished IL-23-induced XBP1s (**Fig. 3B**). Congruously, 4μ8C downregulated the mRNA expression of IRE1-XBP1s downstream genes, *Xbp1s*, *Becn1* (encoding autophagy factor Beclin-1), *Ddit3*, and *Hspα5* ^54–56^ (**Fig. 3C**), and decreased the mRNA and protein levels of TH17 cytokines (**Fig. 3C-D**). Therefore, IRE1 mediates IL-23- induced the expression of XBP1s and cytokines by TH17 cells.

### XBP1s mediates IL-23-driven secretory function of TH17 cells

XBP1s is broadly involved in the secretory function of immune cells ^29, 30^. We have observed above that IL-23 induces the IRE1−XBP1s axis. However, it is not clear whether XBP1s is required for IL-23 to promote the cytokine production by TH17 cells. To explore this, we performed siRNA-mediated *Xbp1* gene silencing in differentiated TH17 cells in the presence or absence of recombinant IL-23. We found that in comparison with scramble siRNA treatment, siXbp1-mediated *Xbp1* knockdown diminished basal and IL-23-induced protein expression of XBP1s (**Fig. 4A**) and mRNA expression of *Xbp1s* and XBP1s targets, *Becn1*, *Hspa5*, and *Ddit3* ^54–56^ (**Fig. 4B**). Consistently, IL-23-induced expression of TH17 cell signature cytokines at mRNA and/or protein levels was reversed by *Xbp1* gene silencing (**Fig. 4B-C**). In summary, IL-23 regulates TH17 cell secretory function in an XBP1s-dependent manner.

**Figure 4.**
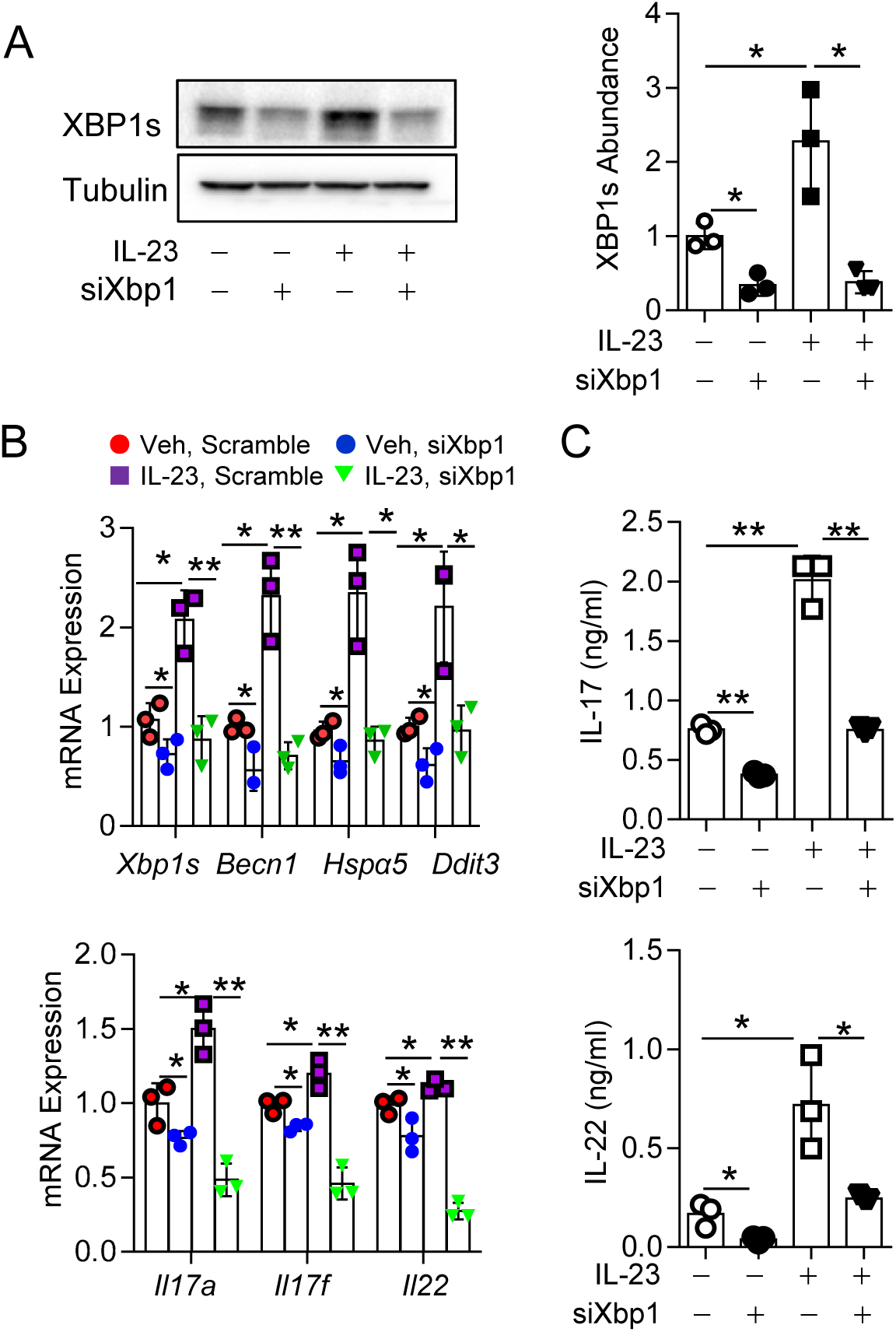
IL-23 promotes the secretory function of TH17 cells via induction of XBP1s. (A) Western blot of XBP1s in TH17 cells treated with or without IL-23 following transfection of siXbp1 or scramble (Sc) siRNA. XBP1s abundances were relative to Tubulin. (B) RT-qPCR of mRNA expression of indicated genes in TH17 cells treated as (A). mRNA abundances were normalized to an internal housekeeping gene *Actb*. (C) ELISA of cytokines, IL-17 and IL-22. Data (means ± SD) were pooled from 3 independent experiments. Student’s *t* test, \**p*< 0.05, \*\**p*< 0.005.

### JAK2 mediates cytokine-driven activation of the IRE1-XBP1s axis in TH17 cells

During UPRs, it is thought that after dissociation from BiP, IRE1 forms homodimer and trans-phosphorylate its partner unit, leading to activation of its RNase and kinase activity ^46, 47^. However, cytokines, leptin and IL-3, induce IRE1 activation in a kinases (MEK and PI3K, respectively)-dependent manner ^41, 44^. These observations prompt us to test whether a cytokine signal-activated kinase can phosphorylate IRE1. IL-23 and IL-6 (and also leptin) use JAK2 to transduce their signals, leading to STAT3 activation ^45^. We asked whether these cytokines also use JAK2 to activate IRE1 in TH17 cells. Because of the importance of the JAK2−STAT3 pathway in IL-23-mediated TH17 effector function ^62, 63^, we treated TH17 cells with IL-23 and found that administration of IL-23 increased the expression of p-JAK2 (**Fig. 5A**) and p-STAT3 (Supplemental **Fig. S1B**), as well as p-IRE1 and XBP1s (**Fig. 5B**). Next, we asked whether JAK2 is required for IL-23 to induce p-IRE1 and XBP1s. Fedratinib (also named TG101348 or SAR302503) is a specific JAK2 inhibitor approved by FDA for myeloproliferative neoplasms ^64^. Remarkably, addition of Fedratinib blocked JAK2 phophorylation and dramatically decreased p-IRE1 and XBP1s expression in TH17 cells; JAK2 inhibition diminished the ability of IL-23 to upregulate p-JAK2, p-IRE1, and XBP1s (**Fig. 5B**). Fedratinib also inhibited the expression of UPR genes and TH17 cytokines in the presence and absence of IL-23 (**Fig. 5C-D**). Accordingly, siRNA-mediated JAK2 gene silencing also diminished the effects of IL-23 on the activation of IRE1 and the induction of XBP1s and TH17 cytokines (Supplemental **Fig. S2**). Taken together, JAK2 is necessary for IL-23 (and likely also IL-6 and leptin) to induce the IRE1-XBP1s axis in TH17 cells.

**Figure 5.**
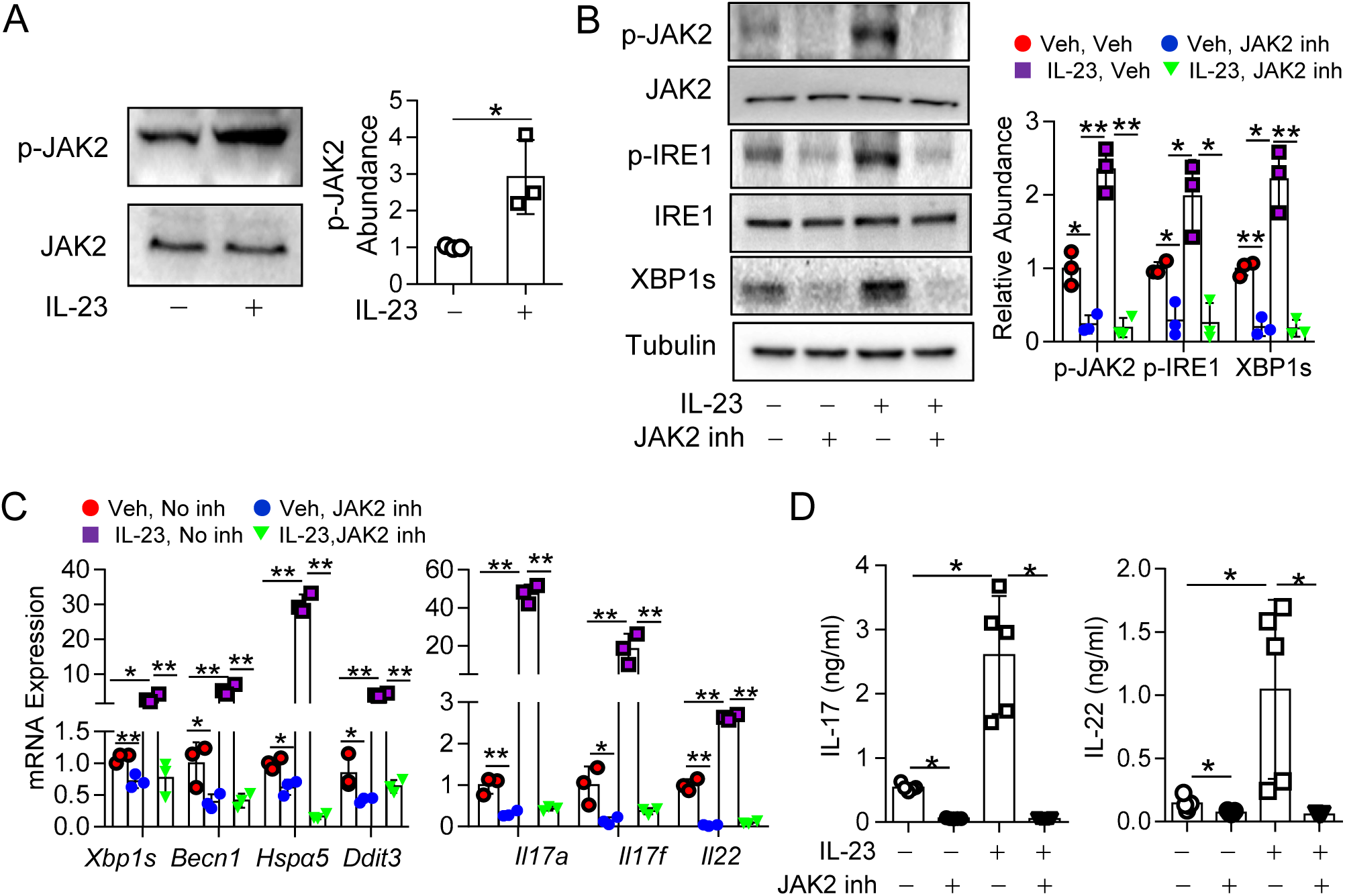
JAK2 is required for IL-23 to induce the IRE1−XBP1s axis in TH17 cells. (A) Western blot of p-JAK2 in TH17 cells treated with or without IL-23. p-JAK2 abundances were relative to JAK2. (B) Western blot of p-JAK2, p-IRE1, and Xbp1s in TH17 cells following JAK2 inhibition (inh) with Fedratinib or a vehicle in the presence or absence of IL-23. Abundances of p-JAK2, p-IRE1, and XBP1s were normalized to JAK2, IRE1 and Tubulin, respectively. (C) RT-qPCR of mRNA expression of indicated genes in TH17 cells treated as (B). mRNA abundances were normalized to *Actb*. (D) ELISA of indicated cytokines in TH17 cell culture supernatant treated as (B). Data (means ± SD) were pooled from 3 (A-C) or 5 (D) independent experiments. Student’s *t* test, \**p* < 0.05, \*\**p* < 0.005.

### p-JAK2 interacts with and directly activates IRE1

We have demonstrated above that IL-23 (and maybe also IL-6 and leptin) signals induce the IRE1-XBP1s pathway via activation of JAK2. To further understand how cytokine-activated JAK2 promotes phosphorylation of IRE1, we first examined whether JAK2 could be physically close to IRE1. *In-vitro* differentiated TH17 cells were starved and reactivated with glass-bound anti-CD3 and anti-CD28 in the presence of IL-23. Following fixation and permeabilization, the cells were labelled with anti-p-IRE1 and anti-p-JAK2. Confocal microscopy revealed that the p-IRE1 stain exhibited perinuclear and perimembranous patterns and p-JAK2 colocalized with p-IRE1 primarily at the perimembranous space with some colocalization also seen in the perinuclear space (**Fig. 6A**). Next, we determined whether p-JAK2 could interact with IRE1. Lysates of IL-23-treated TH17 cells were immunoprecipitated (IP) with anti-p-JAK2, anti-p-IRE1, or isotype control antibodies, and the immune complexes were analyzed by western blot. Anti-p-JAK2, but not an isotype control antibody, co-immunoprecipitated p-IRE1 and p-JAK2 (**Fig. 6B**); consistently, anti-p-IRE1 pulled down p-JAK2 and p-IRE1 (**Fig. 6C**). In vitro, p-JAK2, isolated by IP from IL-23 treated TH17 cells, could activate recombinant IRE1 (**Fig. 6D**). In summary, p-JAK2 colocalizes with and binds to IRE1, and directly phosphorylates IRE1; activated IRE1 subsequently regulates the XBP1s-mediated secretory function of TH17 cells and other functions through its kinase activity (see schematic in **Fig. 6E**). This exogenous kinase activity of JAK2-mediated activation of IRE1 elucidates a previously unexplained phenomenon observed in several early studies including ours ^41, 44^, which is distinct from the traditional view that during ER stress, IRE1 auto-phosphorylation leads to its activation.

**Figure 6.**
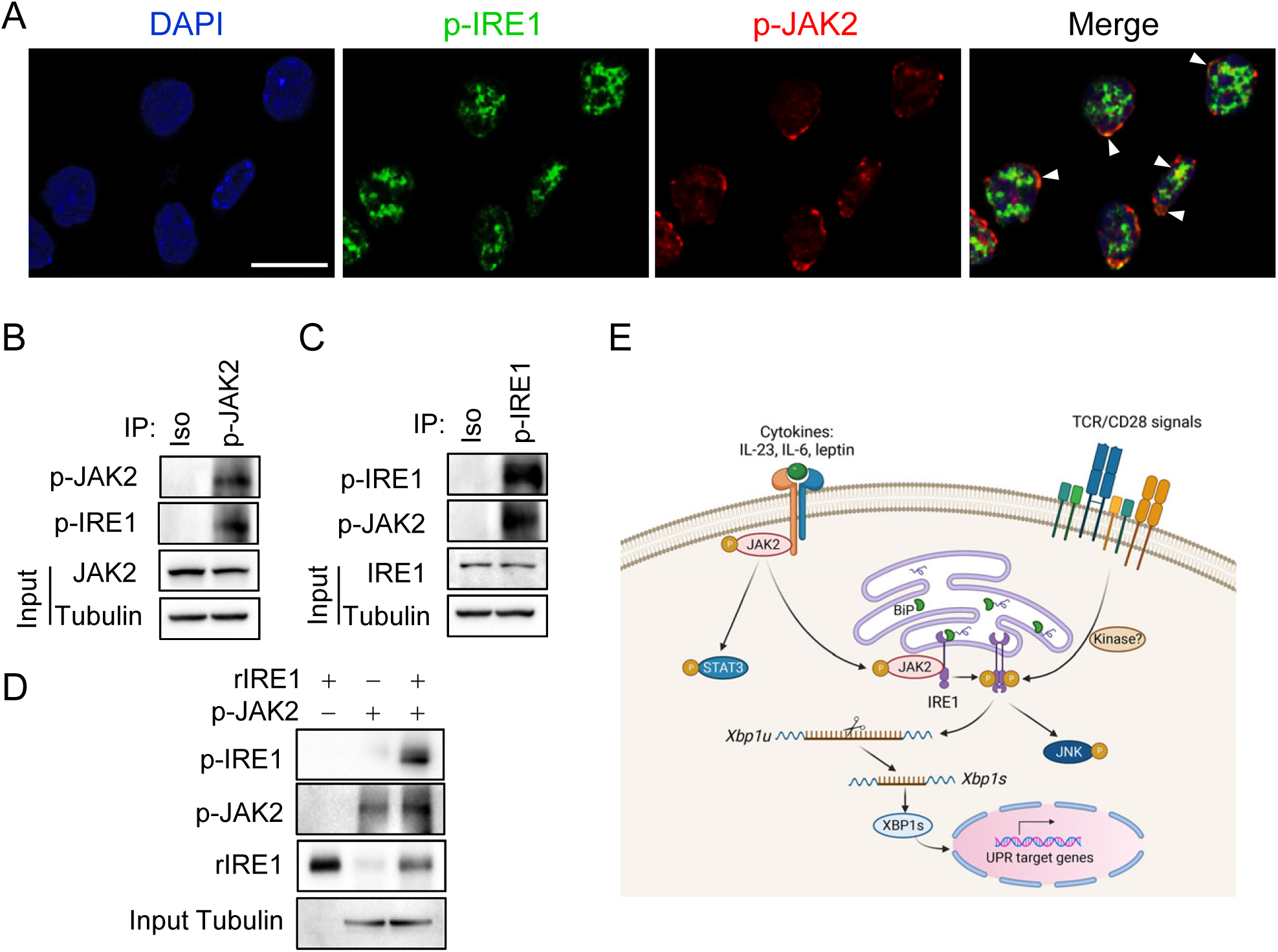
p-JAK2 interacts with and activates IRE1. (A) Immunofluorescence stain and colocalization of p-JAK2 (red) and p-IRE (green). DAPI marked cell nuclei (blue). Arrow heads indicate overlaps of p-JAK2 and p-IRE1. Scale bars, 10 μm. (B-C) Co-immunoprecipitation of p-JAK2 with p-IRE1 by anti-p-JAK2 (B) or anti-p- IRE1 (C). Iso, isotype control antibody. Tubulin and IRE1 or JAK2 were used as a loading control. (D) p-JAK2 phosphorylates recombinant IRE1 (rIRE1) *in vitro*. p-JAK2 was isolated by IP from IL-23 activated TH17 cells and incubated with rIRE1 (shorter than endogenous IRE1). p-IRE1 and inputs p-JAK2 and rIRE1 were detected by western blot. Tubulin was used as a loading control for IP of p-JAK2. Data shown are a representative of 2 (D) or 3 (A-C) experiments. (E) Schematic of extracellular signal-driven activation of the IRE1-XBP1s axis.

### IL-23 induces the JAK2−IRE1−XBP1s axis *in vivo* and enhances fungal asthma

Our above results demonstrate that *in vitro*, IL-23-mediated TH17 cell secretory function is largely dependent on the induction of the JAK2−IRE1−XBP1s axis (**Figs. 3-5**). To understand whether IL-23 also acts on TH17 cells *in vivo* as it does *in vitro* and subsequently drives fungal asthma, we sensitized C57BL/6 (B6) mice i.n. with *C. albicans* extract plus OVA and rechallenged the mice i.n. with OVA only (to avoid the induction of endogenous IL-23 by *C. albicans* ^49, 50^) in the presence or absence of recombinant IL-23. Lung infiltrating CD4^+^IL-23R^+^ TH17 cells from mice receiving IL-23 had increased levels of XBP1s, p-JAK2, and p-IRE1 (**Fig. 7A**) compared with those from the control mice, suggesting that IL-23 activates the JAK2−IRE1−XBP1s pathway *in vivo*. Following activation of the UPR pathway, administration of IL-23 heightened the expression of TH17 cytokines, IL-17 and IL-22, in BALF (**Fig. 7B**) and in lung associated mediastinal lymph node (LLN) cells after *ex vivo* recall with OVA (**Fig. 7C**). IL-23 treatment did not alter the expression of TH2 cytokines, IL-4, IL-5, and IL-13, or TH1 cytokine IFNγ in BALF or LLN recall supernatant (**Fig. S3A-D**). As expected, LLN cells of IL-23-treated animals had increased frequencies and numbers of TH17 cells than those of control mice (**Fig. 7D**). Moreover, we found increased numbers of CD4^+^ T cells and CD4^+^ IL-23R^+^ TH17 cells, accompanying elevated numbers of macrophages and neutrophils, in BALF of mice receiving IL-23 in comparison with the counterparts of the control mice (**Fig. 7E-F**). Histological analyses of lung tissues showed that IL-23 treatment resulted in elevated immune cell infiltration in the alveolar spaces, which contained increased numbers of neutrophils relative to vehicle treatment (**Fig. 7G**). Taken together, IL-23 activates the JAK2− IRE1−XBP1s pathway in TH17 cells *in vivo*, promoting TH17 responses and neutrophilic fungal asthma.

**Figure 7.**
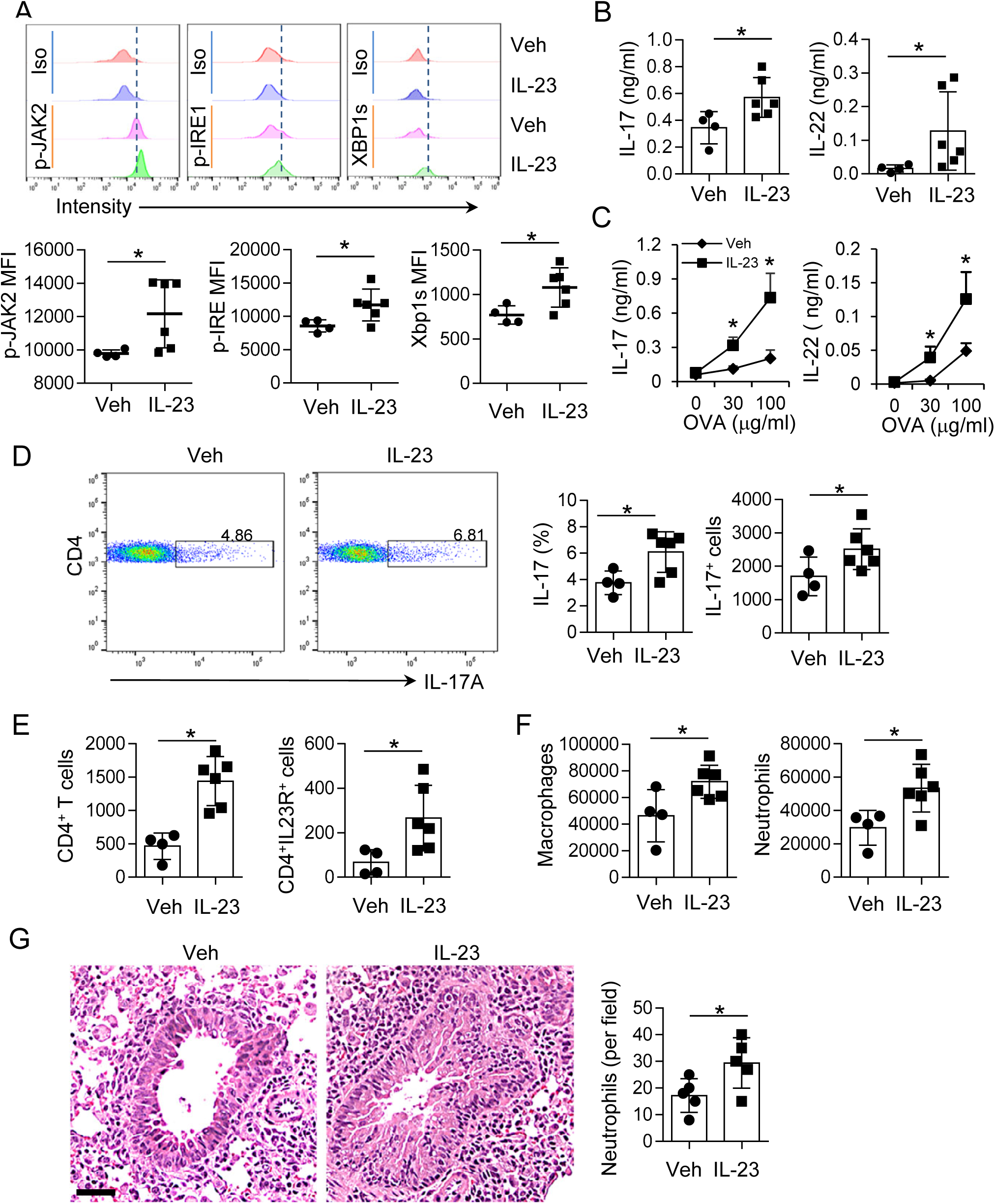
IL-23 activates the JAK2−IRE1−XBP1s pathway *in vivo* and enhances Candida-caused asthma. B6 mice were sensitized with *C. albicans* extract plus OVA followed by rechallenges with OVA in the presence IL-23 or a vehicle. (A) Intracellular stain of XBP1s, pJAK2 and pIRE1 in lung infiltrating TH17 cells on a CD3^+^ CD4^+^ IL-23R^+^ gate. (B-C) ELISA of cytokine expression in BALF (B) and culture supernatant of LLNs after ex vivo recall with OVA (C). (D) Intracellular stain of IL-17-expressing cells in LLNs on a CD3^+^ CD4^+^ gate. (E) Profiles of CD4^+^ T cells and TH17 cells in BALF. (F) Profiles of macrophages and neutrophils in BALF. (G) H&E stain of lung sections. Scale bar, 50µm. Right, neutrophil counts were average numbers per field (x20). Data (mean ± SD) are a representative of two experiments. n = 5 per group. Student’s *t* test, \**p*< 0.05.

## Discussion

ER stress associated UPRs are broadly involved in disease severity and inflammatory phenotype. Studies have revealed that the IRE1-XBP1 pathway plays an essential role in many human pathological conditions, such as inflammation, neurodegenerative diseases, brain and heart ischemia, metabolic disorders, liver dysfunction, and cancers ^65, 66^. Targeting this pathway has emerged as a promising therapeutic intervention against these diseases. Asthma manifests chronic airway inflammation encompassing productive immune infiltration in the airway and the peribronchial and perivascular spaces. GWAS studies have revealed the ER stress factor ORMDL3 as a risk factor of asthma ^32–34^, including severe asthma ^35, 36^. The ORMDL3-ATF6 pathway has been shown to promotes human airway smooth muscle and epithelial cell hyperplasia ^37, 39^. In an *Alternaria* asthma model, ORMDL3 induces ATF6 in airway epithelial cells and in turn promotes T2-high airway disease ^38^. Despite that in airway structural cells, the role of UPRs in TH cells, the driving and maintenance force of chronic airway inflammation, remain poorly understood. Recently, we reported that ATF6 regulates TH2 and TH17 responses and contributes to a mixed granulocytic phenotype of asthma ^42^. However, it is not clear whether IRE1, another effector arm of UPRs, plays a role in asthma, especially neutrophilic asthma. We and others have shown that the IRE1a-XBP1 pathway regulates TH2 responses in T2-high airway inflammation ^40, 41^. In this study, we found that cytokine-driven activation of IRE1 controls the expression of ER stress factors that mediate TH17 cell secretory function and intensify a neutrophilic airway response in Candida-caused asthma model.

UPRs are the central of protein synthesis and secretion. Upon extracellular stimulation, T lymphocytes produce massive amounts of signature cytokines and express increased levels of transcription factors, kinases, signal and structural proteins, and surface markers. Overload of unfolded (or misfolded) proteins in ER leads to ER stress that activates the UPR pathways, IRE1, ATF6, and PERK ^29–31^. Among these, ATF6 is cleaved into an active ATF6 fragment (ATF6f) transcription factor, and ATF6f translocates into the nucleus and transactivates a set of ER stress-related genes, including *Xbp1*. Meanwhile, endoribonuclease IRE1 undergoes phosphorylation and p-IRE1 excises a 26-nucleotide fragment from unspliced *Xbp1* mRNA and forms functional *Xbp1s* mRNA. XBP1s transcribes an array of genes that increase ER capacity, which increases the secretion of a cell, and promote autophagy ^29–31, 67^. The IRE1 and ATF6 pathways together resolve the endogenous cellular stress of unfolded proteins and increase secretory function of the cell, whereas activation of PERK, the third arm of UPRs, leads to induction of autophagy, cell cycle arrest, and even apoptosis if ER stress cannot be resolved. TH17 cells, belonging to a type of secretory lymphocytes, play a critical role in neutrophilic inflammation. However, little is known whether and how UPRs regulate the function of TH17 cells. In this study, we found IL-23, IL-6, and TCR-costimulatory signals can dramatically induce p-IRE and its downstream XBP1s expression in TH17 cells, indicating that extracellular stimuli may influence TH17 function through upregulation of the IRE1−XBP1s axis of UPRs.

How extracellular stimulation, e.g. cytokine signals, activates IRE1 is entirely unclear. The ER transmembrane protein IRE1 contains 3 domains: ER luminal domain, transmembrane domain and cytosolic RNase domain. The luminal domain interacts with ER chaperone BiP maintaining an inactive form of IRE1. Misfolded/unfolded protein binds to BiP leading to its dissociation from IRE1; IRE1 is then proposed to undergo homodimerization and trans-autophosphorylation, leading to conformational change that promotes IRE1 activation ^46, 47^. Although trans-autophosphorylation is generally considered for the activation of IRE1, it may not reflect the situation *in vivo*. Analysis on a luminal domain mutation in yeast IRE1 indicates that BiP dissociation and self-association of IRE1 are not sufficient, but an additional ER-dependent unknown change on its luminal side is required, for IRE1 activation^68^. Furthermore, in pro-B cell Ba/F3, IL-3 induces IRE1 activation and XBP1s expression ^44^. This effect can be blocked by PI3K inhibitor LY294002, suggesting an essential role of the PI3K cascade in IRE1 activation. We have also observed that in TH2 cells, leptin activates IRE1 in a MEK-MAPK- and mTOR-dependent manner ^41^. Therefore, besides the ER-dependent luminal change of IRE1, exogenous kinase activities may act on its cytosol side and contribute to IRE1 phosphorylation. In the current study, we found that cytokines, e.g. IL-23 (and likely also IL-6 and leptin), activate JAK2, and p-JAK2 colocalizes with p-IRE1, visualized by confocal microscopy, and interacts with IRE1, revealed by co-IP assays. Furthermore, p-JAK2 directly phosphorylates recombinant IRE1 *in vitro*. Both pharmaceutical inhibition and siRNA-mediated silence of JAK2 diminish the effects of IL-23 on induction of the IRE1−XBP1s axis. Taken together, p-JAK2 can access and directly activate IRE1. Even though, we always observe basal activation of IRE1 in the absences of any known extracellular stimuli (**Figs. 3, 5, 6**). Therefore, we cannot rule out trans-autophosphorylation as a highly possible mechanism for IRE1 activation under certain circumstances. Nevertheless, the noncanonical activation of IRE1, driven by cytokine signals (and maybe also TCR/costimulatory signals), governs UPRs and mediates the secretory function of TH17 cells.

An array of extracellular stimuli activates the IRE1−XBP1s axis. Though, it is not clear whether these signals behaviour *in vivo* as they do *in vitro*. Of the TH17 inducing factors, IL-23 stimulates the development and maintenance of pathogenic TH17 cells ^25, 57–60^. In steroid-resistant severe asthma, the IL-23−TH17 axis drives neutrophilic airway inflammation ^69, 70^. IL-23 binds the IL-23R−IL-12Rβ1 receptor complex, leading to activation of JAK2 and TYK2 kinases, and subsequently activates STAT3 and STAT4, respectively ^62, 71^. It is well known that IL-23 functions through STAT3 and/or STAT4-mediated transcriptional control, which directs the establishment, maintenance and even phenotype switch of TH17 cells, whereas little is known on its role in posttranscriptional control that determines the altitude of cytokine production, cell size, and cell homeostasis. We found that upregulation of XBP1s is required for IL-23 to induce the expression of TH17 cell signature cytokines as knockdown of *Xbp1* diminishes the effect of IL-23 on the induction of TH17 cell cytokines and the UPR genes. In a fungal asthma model, administration of recombinant IL-23 induces the JAK2−IRE1−XBP1s axis, leading to a hyper-reactive phenotype of TH17 cells characterized by increased production of cytokines and exacerbation of *C. albicans* extract-induced neutrophilic airway inflammation.

In summary, our results indicate that IRE1 is upregulated in asthmatic lungs and in TH17 cells, and noncanonical activation of IRE1 by extracellular stimuli, especially cytokine signals, results in intensive cytokine production by TH17 cells and promotes neutrophilic airway inflammation. These findings may aid to design a novel therapeutic approach for treatment of steroid-resistant severe asthma and other TH17-mediated diseases.

## Materials and Methods

### Mice

All mice used in this study were on the B6 background. *Ern1*^fl/fl;Cd4Cre^ mice were generated by crossing *Ern1*^fl/fl^ with Cd4Cre mice ^52, 53^. All mice were housed in the specific pathogen-free animal facility and animal experiments were performed with protocols approved by the Institutional Animal Care and Use Committee of the University of New Mexico Health Sciences Center.

### Induction and analysis of neutrophilic asthma

6–8-week old sex-matched mice were immunized i.n. with 50 μg *C. albicans* extract (prepared by ultra-sonication from heat-inactivated *C. albicans,* 169246, Greer) and 20 μg chicken ovalbumin (OVA) twice on days 0 and 2, and rechallenged on days 14 and 15. In the IL-23 treatment experiment, the mice were randomly divided into two groups before the rechallenges, one group receiving 20 μg OVA in the presence of 100 ng recombinant IL-23 in 50 µl 0.1% BSA-PBS and another group receiving the vehicle. After the mice were euthanatized by exsanguination under anaesthesia one day post the last dose of allergen, BALF, LLNs and lungs were immediately collected and stored on ice for analysis of infiltrates and immune responses. One lung lobe was fixed in 2% paraformaldehyde for histology and the other lobes were meshed in 1 ml PBS with 120-micron nylon mesh. 5% of lung single cell suspension was removed for profile of CD45^+^ immune infiltrates. The remaining lung suspension was spun to obtain BALF for ELISA measurement of cytokines, and the cell pellet was suspended and treated with a 37% Percoll (17089101, GE Healthcare Life Sciences) density gradient to enrich lymphocytes for flow cytometry assays. Both sexes were used in the experiments, but no significant gender differences were observed.

### TH17 cell differentiation

CD4^+^CD25^−^CD62L^+^ naïve T cells were isolated from splenocytes and lymph node cells of indicated mice and differentiated in a TH17-polarizing condition (2 ng ml^−1^ TGFβ, IL-6 10 ng ml^−1^, 2 μg ml^−1^ anti-IFNγ, and 2 μg ml^−1^ anti-IL4) using plate-bound anti-CD3/anti-CD28.

### IL-23 treatment

*In-vitro* differentiated TH17 cells were rested on a plate without anti- CD3/anti-CD28 for overnight and re-stimulated with plate-bound anti-CD3/anti-CD28 in serum free medium (CTS^TM^ OpTmizer^TM^ T-Cell Expansion SFM, Life Technologies) in the presence or absence of 10 ng ml^−1^ IL-23. The resulting cells were used for indicated assays to measure the expression of cytokine and signal proteins and TH17 signature mRNAs.

### Gene silencing

*In-vitro* differentiated TH17 cells were starved in serum free medium for 24 h. The next day, the cells were transfected with siJak2, siXbp1or a scramble siRNA (40 nm) (Santa Cruz Biotechnology) by electroporation with Neon transfection system (1600v, 20 mS/pulse, 2 pulse; Thermo Fisher Scientific) and incubated on an anti-CD3/anti-CD28 coated plate for 48 h. After confirmation of knockdown of the target gene, the siRNA transfected TH17 cells were treated with or without IL-23 and subjected to different treatments and assays as indicated.

### Treatment with JAK2 and IRE1 inhibitor

After starvation, *in-vitro* differentiated TH17 cells were treated with 2 µM JAK2 inhibitor Fedratinib (TG101348, S2736, Selleckchem), 10 µM IRE1 inhibitor 4μ8C (Cat. #S7272, Selleckchem), or a vehicle for 5 h, following treatment with or without IL-23. The resulting cells were used for indicated measures.

### Flow cytometry

Antibodies against CD4 (RM4.5), CD11b (M1/70), CD11c (N418), Ly-6G (1AB-Ly6g), SiglecF (E50-2440), IL-17 (eBio17B7), and IL-23R (12B2B64) were purchased from eBioscience. Anti-XBP1 (ab220783) and anti-p-IRE1 (S724) (ab48187) were from Abcam, and p-JAK2 (Y1007, Y1008) (44-426G) from Invitrogen. Phospho-protein stain was performed using an optimized protocol as follows: Following stain with antibodies against surface markers CD4 and IL-23R (12B2B64, Biolegend), the cells were fixed by 2% paraformaldehyde for 10 min at room temperature and washed once with cold PBS. The cells were then permeabilized and meanwhile stained with anti-p-JAK2 or anti-p-IRE1 antibody in permeabilization buffer [0.1% Saponin (S7900, Sigma-Aldrich) and 0.1% BSA in PBS] in the presence of Phosphatase Inhibitor Cocktail B (C2118, Santa Cruz Biotechnology) for 50 min at 4 °C with 1-2 agitations. The resulting cells were then stained by a FITC-conjugated anti-rabbit IgG secondary antibody (4300581, eBioscience) and analyzed on an Attune NxT Flow Cytometer (Thermo Fisher Scientific). The data were processed by Flojo software (FlowJo, LLC).

### Western blot and co-IP assays

*In-vitro* differentiated TH17 cells following 24-h starvation were subjected to various treatments as indicated and cell lysates were prepared for immunoblot analysis. For co-IP, the cells were treated with 10 μM MG-132 (S2619, Selleckchem) for 5 h with or without IL-23 treatment for the last 7, 15 or 30 min before lysis. Protein A/G magnetic beads were incubated with 2 μg anti-p-JAK2 (44-426G, Invitrogen), anti-IRE (NB100-2324, Novus), anti-p-IRE1 (ab48187, Abcam) or an isotype control antibody in the presence of Phosphatase Inhibitor Cocktail B for overnight at 4°C. The bead– antibody complexes were washed with cold lysis buffer and then incubated with whole-cell lysates for 1 h at 4°C with agitations. Following washes with cold lysis buffer, the IP complexes were then resuspended in Laemmli’s sample buffer and boiled for 5 min. After removal of magnetic beads by brief spin, the samples were then subjected for immuoblot analysis. Immunoblot antibodies, anti-XBP1 (M186, sc-7160) and anti-JAK2 (C-10, sc- 390539) were purchased from Santa Cruz Biotechnology, anti-α-Tubulin (eBioP4D1) was from eBioscienceand anti-IRE (NB100-2324) was from Novus.

### Kinase assay

p-JAK2 was immunoprecipitated from IL-23-treated TH17 cells as described above. The IP complexes were washed three times with HEPES washing buffer (25 mM HEPES–KOH, 20 mM KCl, pH 7.4) and suspended in 1 × kinase reaction buffer (200 μM ATP, 25 mM Tris, 10 mM MgCl_2_, 2 mM DTT). 100 ng of Recombinant mouse IRE1 protein (ab268540, Abcam) was added into and incubated with the IP product at 30℃ for 30 min. The reaction was terminated by adding 20 μL of 1 × SDS sample loading buffer and denatured at 100°C for 5 min. The resulting samples were analyzed by Western blotting.

### Confocal microscopy

TH17 cells following 5-h starvation were spun on anti-CD3 and anti- CD28-coated glass coverslips in the presence of IL-23 and cultured for 30 min. After fixation with 2% paraformaldehyde and permeabilization with 0.1% Triton X-100 for 15 min at room temperature, the samples were blocked with 1% BSA and labelled with anti-p-IRE1 (ab48187, Abcam) in the presence of Phosphatase Inhibitor Cocktail B for 30 min and stained with FITC-conjugated anti-Rabbit IgG Fab fragment (111-097-003, Jackson ImmunoResearch). After washes and blockade with rabbit serum, the samples were further stained with Alexa Fluor^®^ 647-conjugated anti-p-JAK2 (ab200340, Abcam). DAPI was used to reveal nuclei. Images were acquired on a LSM800 Airyscan inverted confocal microscope equipped with Zen2 software (Zeiss).

### ELISA

The expression of cytokines in culture supernatant and BALF were measured by ELISA using a standard protocol. For *ex vivo* recall, LLN cells (4 × 10^6^ cells ml^−1^) from asthmatic mice were cultured with various concentrations of OVA for 3 days and the supernatant were collected for ELISA. IL-22 was detected by using a Mouse IL-22 ELISA Kit (B284521, Biolegend) following the manufacturer’s instruction.

### RT-quantitative (q) PCR

Gene mRNA expression was determined by using primers in Table S1 with a CFX Connect Real-Time PCR Detection System (Bio-Rad Laboratories) following reverse transcription with SuperScript Reverse Transcriptase (18064022, Thermo Fisher Scientific). Data were normalized to a reference gene *Actb*.

### Statistical analysis

Results were expressed as mean ± SD. Differences between groups were calculated for statistical significance using the unpaired Student’s *t* test. *P* ≤ 0.05 was considered significant.

### Data Availability

All data generated or analyzed during this study are included in this article.

## Acknowledgements

This work was supported in part by NIH grants HL148337, AI116772, and AI142200 (X.O.Y.), and DK110439 (M.L.), and American Diabetes Association Innovative Basic Science Award 1-17-IBS-261 (M.L.). We acknowledge the University of New Mexico Comprehensive Cancer Center Flow Cytometry Facility, Fluorescence Microscopy Shared Resources, and Autophagy, Inflammation and Metabolism in Disease Center Core Facility (supported by NIH CA118100, GM085273, and GM121176, respectively).

## Author Contributions

X.O.Y., M.L., and X.W. designed the study and coordinated the experiments; D.W., and M.A.M. conducted the confocal microscopy experiments; T.I. and A.K. provided and assisted to maintain *Ern1* floxed mice; D.W., X.Z., K.Z., R.W., A.L., and X.O.Y., performed all other experiments; D.W., and X.O.Y. wrote and all authors reviewed the manuscript.

## Competing Interests

The authors declare that they have no competing interests.

---

**Table S1.**
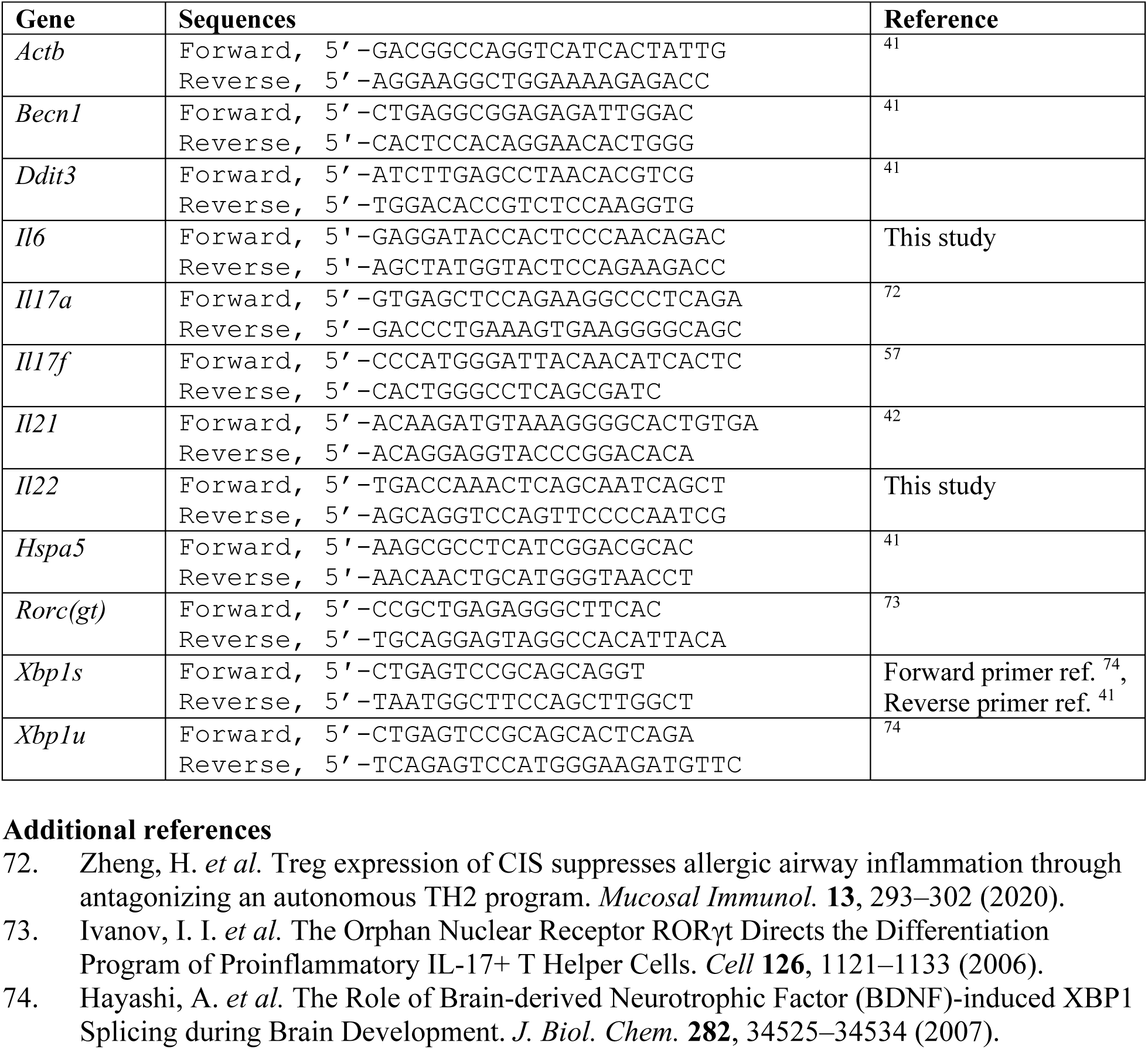
RT-qPCR primers.

**Fig. S1.**
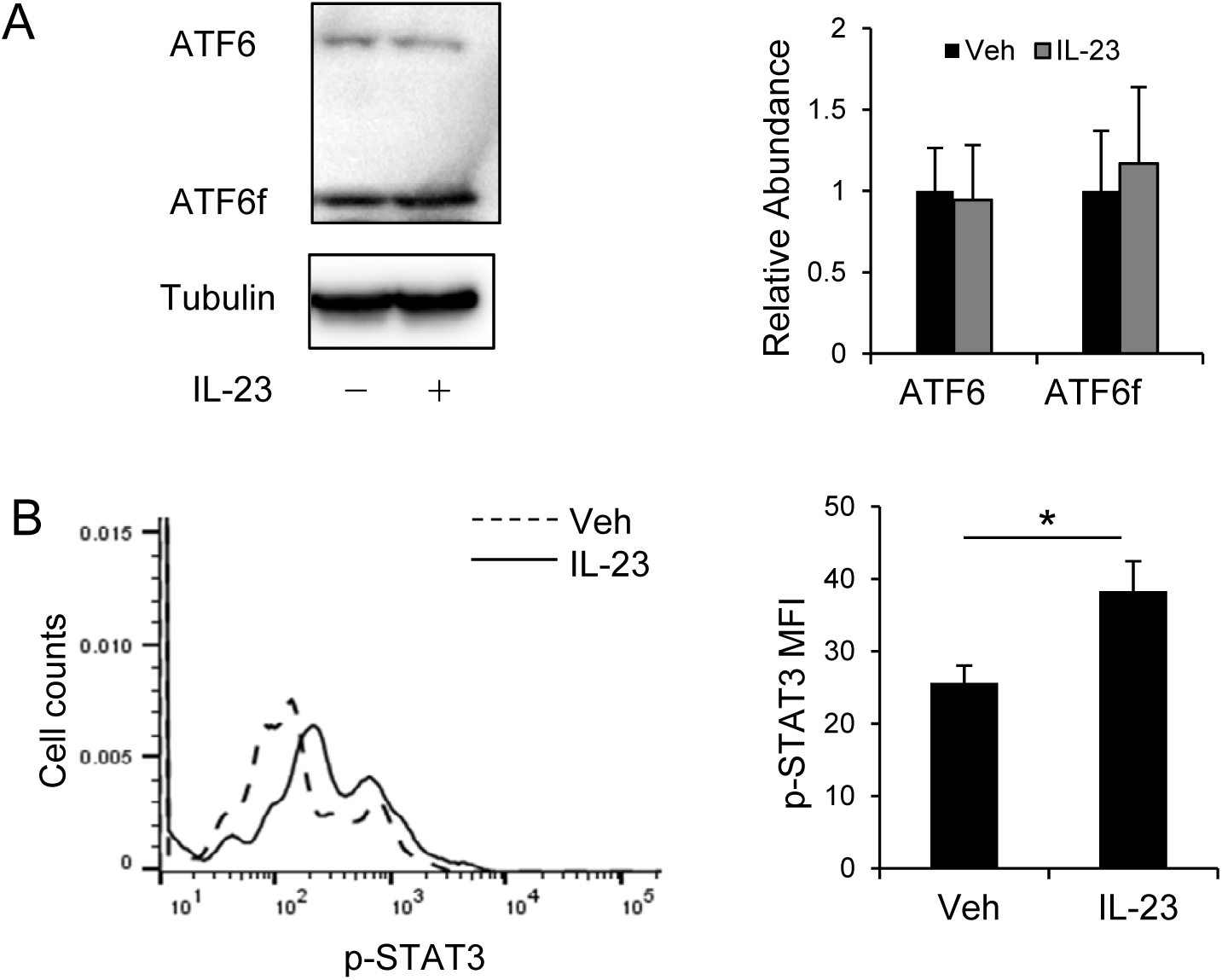
Effects of IL-23 on the activation of ATF6 and STAT3. (A) Western blot of ATF6 and ATF6f (active form of ATF6) in TH17 cells with or without treatment of IL-23. Abundances of ATF6 and ATF6f were normalized to Tubulin. (B) Flow cytometry of p-STAT3 expression in TH17 cells with or without treatment of IL-23. MFI, mean fluorescence index. Data were pooled from 3 individual experiments. \**p* < 0.05.

**Fig. S2.**
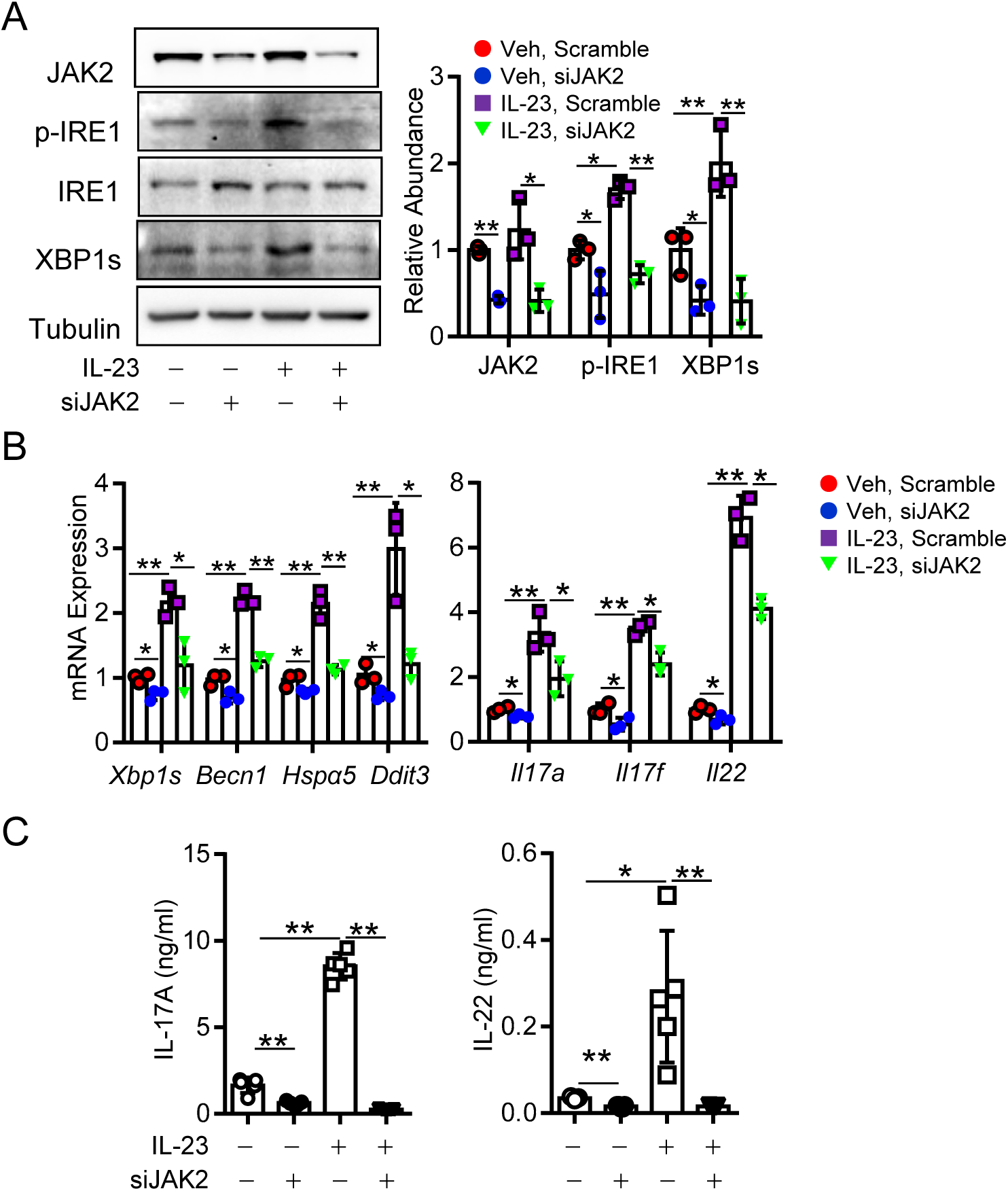
Knockdown of JAK2 diminishes the induction of the p-IRE1-XBP1s axis by IL-23 in TH17 cells. (A) Western blot of p-JAK2, p-IRE1, and XBP1s in TH17 cells treated with or without IL-23 following transfection of siJAK2 or scramble siRNA. Abundances of p-JAK2, p-IRE1, and XBP1s were relative to JAK2, IRE1, and Tubulin, respectively. (B) RT-qPCR of mRNA expression of indicated genes in TH17 cells treated as (A). mRNA abundances were normalized to *Actb*. (C) ELISA of indicated cytokines. Data, means ± SD, are a combination of 3 (A, B) or 5 (C) independent experiments. Student’s *t* test, \**p* < 0.05, \*\**p* < 0.005.

**Fig. S3.**
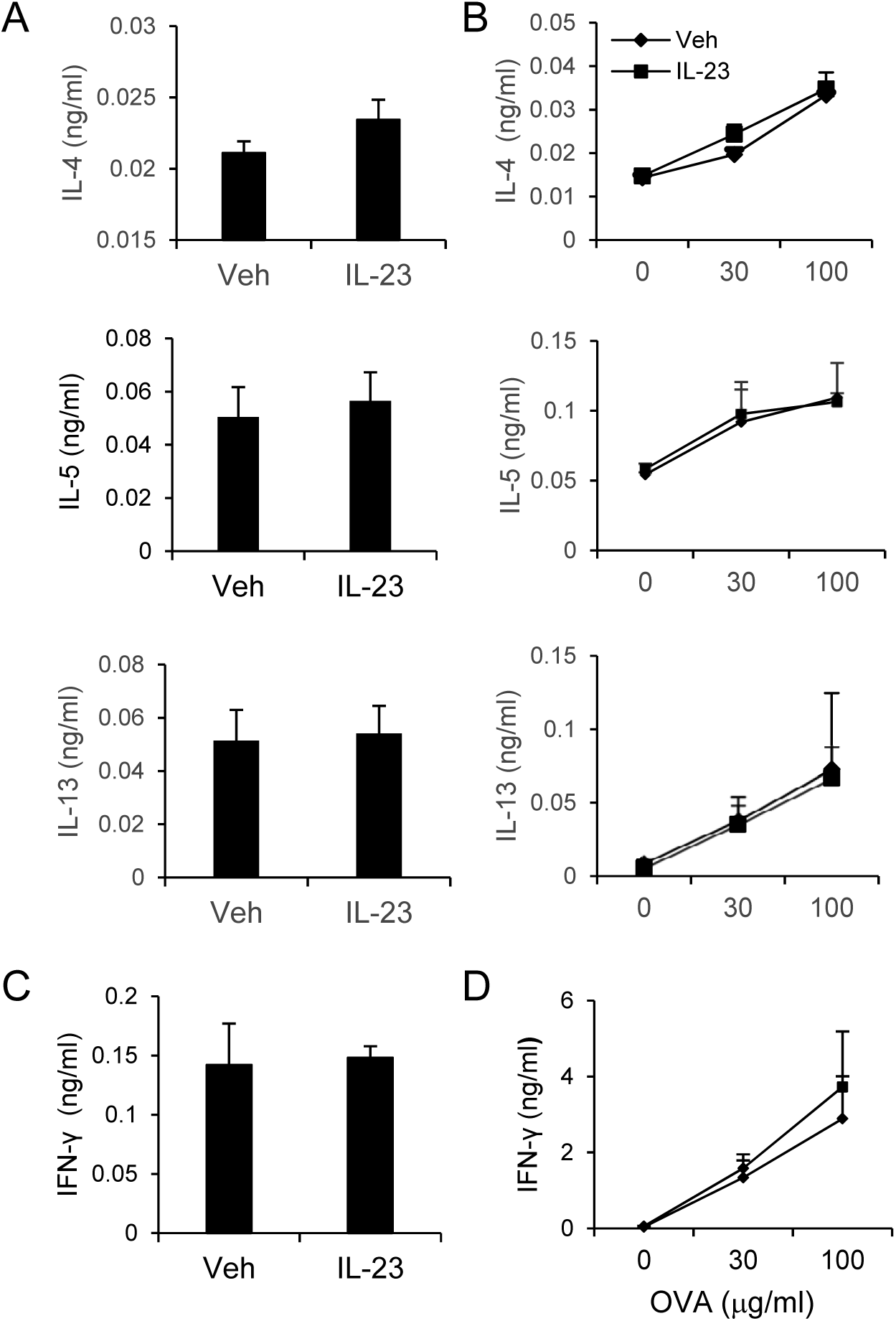
Effects of IL-23 on the expression of TH1 and TH2 cytokines in asthmatic mice. (A-B) ELISA of TH2 cytokines, IL-4, IL-5, and IL-13 in BALFs (A) and culture supernatants of LLNs after ex vivo recall with OVA at indicated concentrations (B). (C-D) ELISA of TH1 cytokine IFNγ in BALFs (C) and culture supernatants of LLNs after ex vivo recall with OVA at indicated concentrations (D). Data are a representative of two experiments (4–6 mice per group). Student’s *t* test, not significant.

## Notes

### Competing Interest Statement

The authors have declared no competing interest.

### Summary of Updates

We fixed the omission of the author list and made minor revision in the text.

